# Differential correlation across subpopulations of single cells in subtypes of acute myeloid leukemia

**DOI:** 10.1101/2022.03.07.483400

**Authors:** Reginald L. McGee, Gregory K. Behbehani, Kevin R. Coombes

**Affiliations:** College of the Holy Cross, Worcester, Massachusetts, United States of America; Department of Hematology, The Ohio State University Wexner Medical Center, Columbus, Ohio, United States of America; Department of Biomedical Informatics, The Ohio State University Wexner Medical Center, Columbus, Ohio, United States of America

## Abstract

Mass cytometers can record 40-50 parameters per single cell for millions of cells in a sample, and in particular, for leukemic cells. Many methods have been developed to cluster phenotypically similar cells within cytometry data, but there are fewer methods to visualize activity and interactions of pairs of proteins across these populations. We have developed a workflow for analyzing correlations associated with predfined populations. By clustering blood samples from acute myeloid leukemia (AML) patients and normal controls using an established algorithm, we obtained a minimum spanning tree of clusters of single cells. Using surface marker expression, we identified clusters on the tree that belonged to phenotypes of interest. Next, we computed correlations between pairs of proteins in each cluster. We developed a novel, coherent, probability-based statistic to test differences between vectors of correlation coefficients. By comparing all combinations of the normal controls under the statistic, we created an empirical distribution that provided a conservative measure of differential correlation. Using this empirically-derived distribution to define significance, we compared pooled samples from AML subtypes and normal controls to detect differential correlations. Given the structure present within this cytometry data set, we found it natural to consider correlations in this manner versus aggregating all data and computing a single correlation. Our results have the advantage that we can localize the statistical measure to determine contributions from particular phenotypic populations. Differentially correlated pairs of proteins can be further explored by considering a population’s distribution of correlation coefficients or biaxially plotting protein expressions within individual cells in a given population. Our approach leads to a better understanding of the nonlinear relationships that exist in the cytometry data.

**Author summary:** We introduce a novel method for analyzing the abundance of single cell data collected by high-throughput technologies. Due to the high dimensionality of such datasets, there is a need for methods to identify significant interactions between genes or proteins. In particular, we are interested in statistical differences between correlations of proteins within populations of cells determined by traditional immunophenotyping techniques. In this paper, we have demonstrated the utility of this new framework in the case of blood samples from individuals with different subtypes of leukemia and compare to healthy controls. The motivation for this application is that differences between can illuminate potential drivers of the disease. We have illustrated an example of how intracellular events can be detected by our statistic. Finally, this method is flexible in that it can also be applied in a variety of contexts where there are vectors of correlation coefficients that need to be compared.

## Introduction

Single cell technologies provide promise for gaining new insights into genetic diseases such as cancer, but the increasing dimensionality of these datasets makes analysis more complicated and the determination of mechanisms that promote the disease less obvious. Mass cytometry records expression levels for tens of proteins on hundreds of thousands of individual cells in collected samples [1]. Moreover, there is additional structure to the data in that each cell has a phenotype, and the samples can be collected from individuals who may have different subtypes of a disease of interest. Investigating high dimensional datasets such as these requires the development of new methodologies, particularly methods that consider the varying levels of structure inherent to the dataset.

Acute myeloid leukemia (AML) is a genetic disease marked by an accumulation of immature myeloid cells in the bone marrow and eventually within the peripheral blood. For individuals diagnosed with AML, the 5-year relative survival rate is 28.5% [2]. The survival rates differ across subtypes of AML and thus understanding genetic and molecular differences between subtypes is of the utmost importance.

Classifying a subtype of cancer requires consideration of factors such of cytogenetics, molecular mutations, and the maturation and phenotype of the cancerous cells. A complication in developing treatments for AML is the heterogeneity observed in these factors. The disease can afflict a variety of cell populations and these populations can play different roles in the diseases. Two known cell populations are leukemic stem cells (LSCs) known to contribute the initialization and recurrence of the disease [3] and leukemic regenerating cells (LRCs) now known to promote disease recurrence after chemotherapy [4]. Thus, metrics for quantifying the activity of cell populations in samples from different disease types, samples from individuals receiving therapy, and samples from the normal population could be useful for clinicians.

Specific karyotypes and mutations are often associated with different blood cancer subtypes. One avenue for attempting to understand the functions of these mutations is considering how protein interactions within cancerous phenotypes from normally observed interactions. Historically, as sequencing data became more ubiquitous, new techniques for data analysis were necessary to determine co-regulated genes and to better understand prognostic factors [5, 6]. It is natural to develop similar techniques as high-dimensional cytometry data continues to become more prominent. We seek to develop techniques for detecting differences in protein co-regulation across both patients and cellular phenotypes.

An advantage of using mass cytometry when studying AML is that it can capture biological phenomena at the level of single cells and is particularly suited to the analysis of complex cell populations. Gating strategies, the consideration and comparison of different parameters collected during a cytometry panel, can be used to identify phenotypes of the cells present in a sample and then to visualize them in two dimensional plots. The identification of phenotypes can be aided by recent techniques for analyzing cytometry data; these range from projections from higher dimensional spaces, where the data lives, into two and three dimensions [7–9] and clustering algorithms that produce trees of clusters where the branches are phenotypically related [10–13]. In both cases, the expression of surface proteins one at a time can help to gate the projected or clustered data. Moreover, there has been work considering differential abundance of cell populations in mass cytometry data [14] given that determining the type of blood cancer and rate of progression depends on the phenotype most closely related to the malignant cells that have accumulated.

In addition to differential abundance of cell populations, differential expression of intracellular proteins is also of interest. A visual analysis of one intracellular protein at a time can be conducted after the clustering or projection methods above have been applied. These approaches can give more prognostic insight into biological features within specific phenotypes, but do not provide information on co-regulation since they only consider one protein at a time. Thus, there is a need for tools that consider more than one protein at a time. In the case of two proteins, determining co-regulation starts with finding positive correlations between proteins. Given that multiple cell populations can be identified within mass cytometry data, co-regulation between two proteins could be determined either for an entire sample or for specific cell populations. Our approach analyzes pairs of proteins across samples that can also be localized to specific cell populations of interest within each sample. The advantage here is that the approach can be applied to other single cell technologies and, more generally, to any structured collection of correlation coefficients.

In this manuscript, we present a technique for determining when protein interactions change in aggregate between samples being compared. That is, we consider when there is a significant change in protein correlations within cells of the observed phenotypes between two samples being compared. We were particularly interested when there are significant differences between samples from patients diagnosed with distinct subtypes of AML as this could point towards underlying mechanisms driving the disease. We begin by describing the AML data used in this study and discuss how it was processed as well as the cell count variability across samples and across subtypes. Next we give a brief description of the established agglomerative clustering algorithm applied to all samples aggregated together; emphasizing that the surface marker expression from each sample was passed to the algorithm to produce a consistent minimum spanning tree of clusters of cells that are phenotypically similar. Next we discuss how the clustered data was processed and begin the derivation of the statistic we use to test differential correlation between samples. In the third section, we have presented results of applying the derived statistic and use traditional gating techniques to investigate the significant changes in correlation detected for particular phenotypes. Finally, we discuss the informatic and translational impact of the findings.

## Materials and methods

### Data description

We investigate existing acute myeloid leukemia data (AML) for intracellular correlations that undergo statistically significant changes when comparing different subtypes of the disease.

The mass cytometry data, originally collected by Behbehani *et al*. [3], consisted of 60 marker readings for 58 samples in ten groups, with eight groups being different AML subtypes. The original study sought to determine aberrant marker expressions in the AML subtype samples when compared to normal controls. Aberrance was determined by a 2-fold difference outside the total variable of the normal samples one marker at a time.

The subtypes considered, as shown in Table 1, were: core-binding factor AML (5 samples), t(8;21) or inv(16) karyotype (4 samples), normal karyotype with a FLT3-ITD mutation (11 samples), normal karyotype with wildtype FLT3 (5), adverse risk karyotype (7 samples), normal karyotype with FLT3-TKD mutation (4 samples). In addition to these groups, there was also a group of fourteen normal control samples and a group of six samples that responded positively to chemotherapy. In all subsequent plots, the subgroup colors were chosen with the Polychrome R package [15].

**Table 1.** Summary of the different subtypes of acute myeloid leukemia (AML) considered in the study by Behbehani *et al*. [3] and reanalyzed here. Shortnames for the different subgroups, the number of samples, and associated colors to help distinguish subgroups are given. The subtypes are compared to 14 samples from healthy donors (associated color: 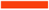) in this study.

| Shortname | Full diagnosis | # of Samples | Color |
| --- | --- | --- | --- |
| CBF | Core-binding factor AML | 5 | Green |
| APL | t(8;21) or inv(16) karyotype | 4 | Orange |
| FLT3-ITD | Normal karyotype with a FLT3-ITD mutation | 11 | Purple |
| FLT3wt | Normal karyotype with wildtype FLT3 | 5 | Pink |
| ARK | Adverse risk karyotype | 7 | Dark Blue |
| FLT3-TKD | Normal karyotype with a FLT3-TKD mutation | 4 | Brown |
| CR | Patients without AML after chemotherapy | 6 | Blue |

There were 53 markers measured that were split across two cytometry panels, each panel having a total of 47 markers (Figure 1). There were 21 surface markers as well as five intracellular markers that were consistent across the two panels. Each panel had 13 unique intracellular markers corresponding to cell cycle or signaling proteins. The other markers were used for barcoding or were not appropriate to be considered in a correlation study. Thus, a total of 18 markers will be used for our analysis.

**Fig 1.**
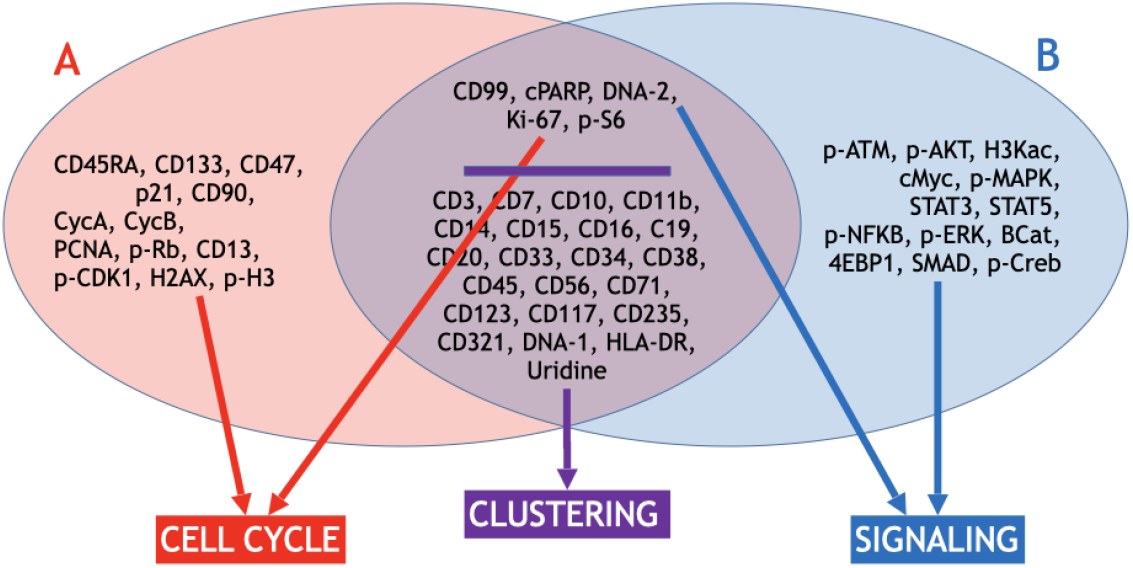
Summary of the biomarkers measured in the mass cytometry data collected in the study Behbehani *et al*. [3]. Two red and blue ovals represent collections of biomarkers (panels) to measure, with the panels corresponding to either cell cycle or intracellular signaling dynamics. There are common biomarkers (purple) between the panels, these are largely surface markers used to cluster individual cells by their phenotype. Differential correlation analysis was performed with the markers distinct to the cell cycle or cell signaling panel (red or blue, respectively).

### Workflow

An overview of the workflow to identify differentially correlated vectors is shown in Figure 2.

**Fig 2.**
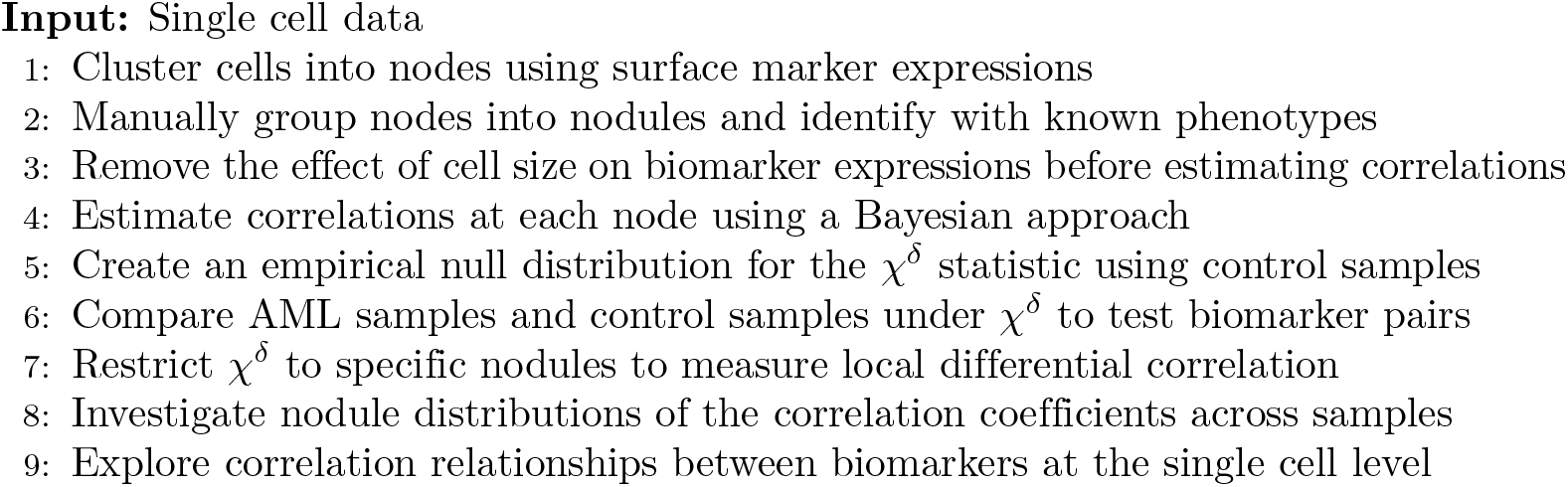
Workflow for identifying differentially correlated biomarkers pairs. The global differential correlation measure can be restricted to a particular nodule to get a local measure of differential correlation. Local investigation and exploration of differential correlation in nodules of interest can be accomplished with bean and biaxial plots. Localization of the statistical measure helps to put the correlation relationships into specific contexts.

### Clustering with SPADE

Spanning tree Progression of Density normalized Events (SPADE) [11] is an approach to extract information about hematopoetic development from mass cytometry data. SPADE uses agglomerative clustering and downsamples dense regions of cells before clustering to ensure rare cell types are not missed. After connecting the nodes into a minimum spanning tree the regions are upsampled and cells are placed into appropriate nodes.

The dataset we consider was clustered in R with SPADE version 1.18.2, using 19 of the 21 surface markers common in the two panels. During a previous analysis with this dataset [3], the surface marker CD56 was omitted because of dimness of the reading and CD99 was used here during analysis as it has been previously used to correlate with disease. We sought a consistent minimum spanning tree for all patients and thus the 116 files were clustered together. 483 nodes was the number of nodes specified to SPADE; the number 483 is chosen due to it being the square root of the average number of cells per sample [25].

More important than a precise number of nodes created by the algorithm, are the phenotypes of the cells within those nodes. Minimum spanning trees produced by SPADE have the advantage that branches can be manually reconciled with known cell phenotypes. We define nodules as collections of nodes that can be identified with a phenotype. The nodules were determined manually by considering combinations of surface marker expressions, gates, from the samples from healthy donors and then the normal cell phenotype groupings determined by the gates were projected onto the tree throughout the paper (Figure 3).

**Fig 3.**
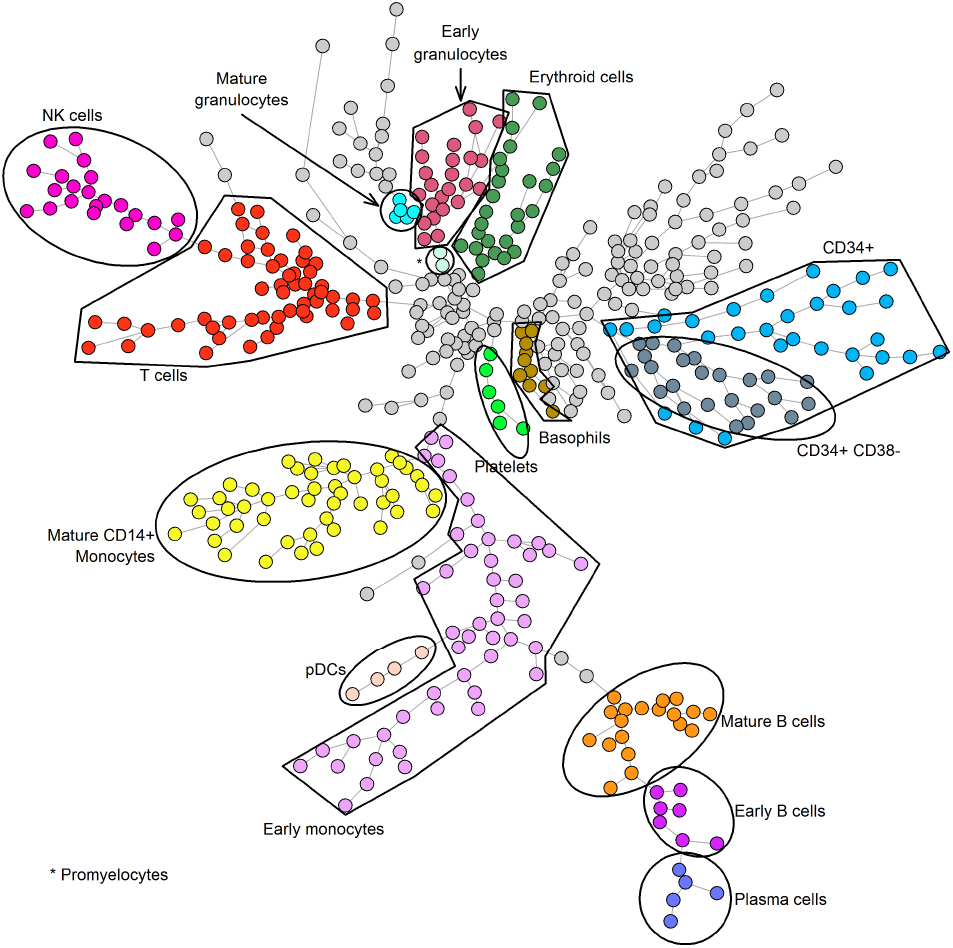
To ensure consistent comparisons between samples during our analysis, a single tree was created using surface marker expressions, shown in Fig 1, from all of the AML samples and samples from healthy donors. The tree was produced by Spanning tree Progression of Density normalized Events (SPADE) [11] considering all samples together, the nodes thus have consistent surface marker expression ranges across all samples. Nodes were reconciled with phenotypes for samples from healthy donors and this gives a sense of the cell phenotypes in the AML samples. There are 16 phenotypes annotated on the plot above and colored to be further distinguished. The remaining cells, left grey, could be classified, but the groups shown were those most of interest for this study.

### Processing clustered data

At each node in the minimum spanning tree, we estimated correlation matrices for the 18 markers to be analyzed at each of the nodes through a Bayesian approach [16]. The estimation circumvents issues that arise when a node is empty or contains only a few cells and attempts to correlate data with built-in R functions. When there are no cells (*N* = 0) in a given node, the method estimates the 18×18 correlation matrix *C*_*N*_ for that node to be the identity matrix *I*_18_. Assuming a Wishart prior distribution, the correlation matrix *C*_*N*_ were estimated as follows when there are *N* cells in a node:

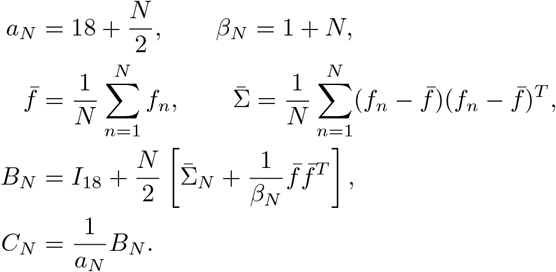

Thus, when there are many cells in a node, then the covariance estimate in *B*_*N*_ overwhelms the prior *I*_18_. Correlation matrices are estimated at every node, and stored in the folders named according to that node, for individual samples, the all-sample pool, and each subtype pool.

### Identifying statistically significant changes in biomarker pairs

In order to find statistically significant changes between the group of normal control samples and groups of samples with different subtypes of AML, we sought a distribution on the Pearson correlation coefficients and created a test statistic *χ*^*δ*^ (Eq 2) related to a Chi Square, *χ*^2^, statistic. Given a marker pair *x* and *y* of the 18 markers in the panel reserved for analysis, there is a correlation coefficient at each of the 483 nodes for each group of samples. When comparing all possible pairs, this resulted in 153 vectors 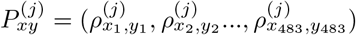, where *j* is a group pooled by sample subtype and each component is a correlation coefficient between markers *x* and *y* in the *i*^*th*^ node determined by SPADE.

Our null hypothesis was that the correlations at node *i* arise from the same distribution and thus their distribution parameters were equal. A Fisher z-transform 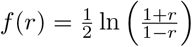 was applied componentwise to 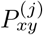 for each *j*. This transform maps the components from a probability distribution on [−1, 1] to a normal distribution with mean 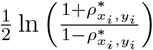 and standard deviation 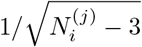, where 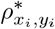 is the true correlation coefficient and 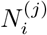 is the number of cells used to estimate 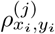. When considering the sum of the 483 squared standardized normal variables

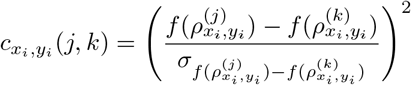

we determined that our statistic for significant aggregate differences in correlation should, in theory, arise from a Chi Square distribution with 483 degrees of freedom, denoted 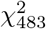.

Due to the number of cells in certain nodes being in the thousands, we regularized our statistic by adding an exchangeability factor *s*_0_ to the standard deviations in 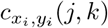 at each node *i*; an idea borrowed from Tusher *et al*. [17]. The value of *s*_0_, shown in Table 6, are uniform for all sample comparisons for either cell signaling or cell cycle data and was determined by minimizing the deviation between 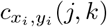 and the expected value of 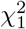.

Intuitively, the addition of the *s*_0_ term serves to prevent oversensitivity at nodes with a large number of cells by preventing the denominator from becoming too small and makes the statistic more conservative overall. Thus, we define our test statistic to be

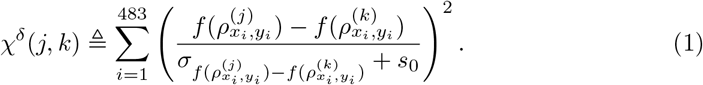

To proceed with hypothesis testing, we chose to define an empirical null distribution using the *χ*^*δ*^ statistic to compare the fourteen normal samples. The motivating idea was that an empirical distribution defined from normal comparisons would provide a baseline for significant differences one could expect when comparing vectors of cytometric correlations and anything beyond this baseline difference could confidently be considered significant.

Using the empirical distribution, we considered nominal *p* values of 0.1, 0.05, 0.01, and 0.001. When considering the nominal *p* value of 0.001 we found the false discovery rate (FDR) value of 0.117 and 0.030 to be satisfactory for the cell cycle and cell signaling datasets, respectively. A summary of the number of significant findings and FDR values at each nominal p value can be seen in Table 3. In the subsequent analysis, we choose the level *p* = 0.001 for significance.

**Table 2.** Exchangeability factors to minimize the difference between the means of the summands 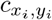 and 1, the mean of a 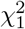. The minimization was performed with the one dimensional optimization function `optimize` in R.

| Cell cycle $s_0$ | Signaling $s_0$ |
| --- | --- |
| 0.02188310 | 0.02092582 |

**Table 3.** Summary of the number of biomarker pairs found to globally differentially correlated, across all comparisons of subtype groups to normal controls, and the false discovery rates (FDR) at different p values for each dataset. Significance levels were determined using an empirical distribution derived from comparing the normal samples under the *χ*^*δ*^statistic. For subsequent analysis we chose the significance level *p* = 0.001.

| Nominal p | Cell cycle;n | Cell cycle;FDR | Signaling;n | Signaling;FDR |
| --- | --- | --- | --- | --- |
| 0.100 | 704 | 0.617 | 837 | 0.519 |
| 0.050 | 508 | 0.428 | 701 | 0.310 |
| 0.010 | 196 | 0.222 | 421 | 0.103 |
| 0.005 | 91 | 0.239 | 308 | 0.071 |
| 0.001 | 37 | 0.117 | 147 | 0.030 |

## Results

We began by using SPADE to produce a minimum spanning tree that incorporated information about the surface markers from all samples in both the cell cycle and signaling experiments (Figure 3). Using the surface marker expression from the fourteen normal control samples, sixteen immunophenotypes of interest were manually reconciled with the nodes determined by SPADE. This reconciliation creates our immunophenotype nodules that are annotated atop the minimum spanning trees. For the purposes of this analysis, we refer to the unannotated cells on the diagram as the nodule of “other cells”. Moreover, the *χ*^*δ*^ statistic can now be restricted to nodes from a particular immunophenotype. For the rest of the paper, the global *χ*^*δ*^ statistic considers all 483 nodes and local *χ*^*δ*^ statistics consider nodes corresponding to particular phenotype nodules. To analyze a differentially correlated pair locally with respect to an immunophenotype nodule, we provide both biaxial plots of marker expressions of individual cells and the distribution of correlation coefficients.

### Adjusting for cell size

In general, larger cells are expected to have higher levels of most markers. So, before the main analysis, we normalized the data to adjust for the effect of cell size, which was measured by an iridium-based cell (IBC) marker. Specifically; we removed the the IBC marker’s contribution to the expression of each feature by performing a linear regression between that feature and the IBC marker. All further analyses were performed on the residuals from these linear regressions.

Shown in Fig 4 are minimum spanning trees before and after the cell size adjustment for each sample at each node. These trees have been colored by the amount of correlation between a particular biomarker pair, histone 3 lysine acetylation (H3Kac) and phosphorylated mitogen-activated protein kinase (pMAPK) whencomparing normal karyotype with FLT3-TKD mutation AML samples to normal controls). This pair of markers was found to be differentially correlated before the cell sizeadjustment but was not determined to be differentially correlated after adjustment. In total, 137 marker pairs were determined to be differentially correlated before adjusting for cell size but not after adjustment. The adjustment overall reduces correlation between the markers and thus further increases the conservativeness of our results. For the rest of the paper, the data presented will have the per-sample-per-node cell size adjustment.

**Fig 4.**
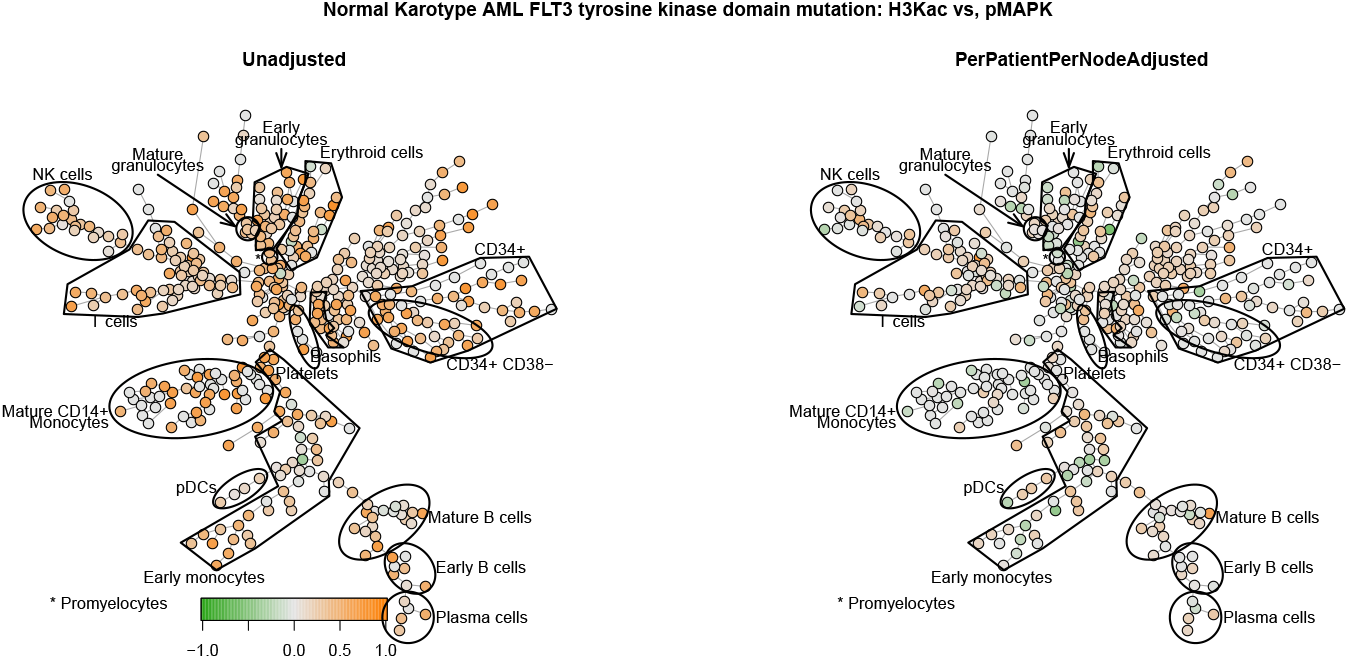
Illustration, on an absolute scale, of the how the adjustment for cell size at every node for each sample removes artificial positive correlation. The minimum spanning trees are colored by correlation coefficients between histone 3 lysine acetylation (H3Kac) and phosporylated MAPK (pMAPK) on a relative scale. The removal was motivated by the expectation that larger cells would have higher expressions of the measured markers. On the left, the correlations are computed with unadjusted data and, on the right, the correlations are computed with residuals from the linear regression of each marker H3Kac and pMAPK on the contribution of the IBC marker (which measures cell size) for every sample at each node. Before the adjustment, the pair H3Kac and pMAPK were found to be globally differentially correlated. Overall, there is a reduction of positive correlation across the minimum spanning tree.

### Statistically significant changes in correlation

We computed *χ*^*δ*^ statistics to test for changes in correlation vectors for each of the 153 possible biomarker pairs in each of the seven groupings of AML samples compared to normal control samples, leading to a total of 1071 comparisons. At the significance level *p* = 0.001, using the cutoffs from the empiorical null distribution (3040 and 1323, respectively), we found 37 significant pairs in cell cycle data and 147 significant pairs in cell signaling data. For both datasets, the normal karyotype with a FLT3-ITD mutation subtype had the most significant pair findings (cell cycle: 10; cell signaling: 35). Some of the differentially correlated pairs served as positive controls, some corresponded to known biological relationships, and some were potentially novel relationships. In the subsequent analysis we will only report on the cell signaling data. The potentially novel biomarker pairs and the groups for which they were found to be differentially correlated compared to the samples from healthy donors are summarized in Table 4. How these global differential correlations results present locally, within the different cell phenotypes within the data, can be explored by plotting the correlation coefficients of the biomarker pairs atop the minimum spanning tree. We illustrate an example of this exploration in Fig 5.

**Table 4.** Summary of potentially novel biomarker pairs and the sample groups for which the pairs were found to be globally differentially correlated compared to normal control samples. The 147 biomarker pairs listed in Table 3 were divided into the categories: positive controls, known biological relationships, and potentially novel relationships.

|  | CBF | APL | ITD | FLT3wt | ARK | TKD | CR |
| --- | --- | --- | --- | --- | --- | --- | --- |
| pS6;H3Kac | No | No | Yes | Yes | No | No | No |
| pS6;CD99 | No | No | No | No | No | Yes | No |
| pATM;H3Kac | Yes | No | Yes | Yes | Yes | Yes | Yes |
| pATM;CD99 | Yes | No | Yes | No | Yes | No | No |
| H3Kac;pMAPK | Yes | No | Yes | Yes | Yes | No | No |
| H3Kac;STAT5 | No | No | Yes | No | No | No | No |
| H3Kac;Ki-67 | Yes | Yes | Yes | Yes | Yes | Yes | Yes |
| H3Kac;CD99 | Yes | Yes | Yes | Yes | Yes | Yes | Yes |
| H3Kac;4EBP1 | Yes | No | Yes | Yes | Yes | Yes | No |
| H3Kac;pCreb | Yes | No | Yes | Yes | Yes | Yes | No |
| pMAPK;CD99 | Yes | No | Yes | Yes | Yes | No | No |
| STAT3;CD99 | Yes | No | Yes | No | No | No | No |
| STAT5;CD99 | No | No | Yes | No | No | No | No |
| Ki-67;CD99 | Yes | Yes | Yes | Yes | Yes | Yes | No |
| CD99;pErk | Yes | No | Yes | No | No | Yes | No |
| CD99;4EBP1 | Yes | Yes | Yes | Yes | Yes | Yes | No |
| CD99;pCreb | Yes | Yes | Yes | Yes | Yes | Yes | No |

**Fig 5.**
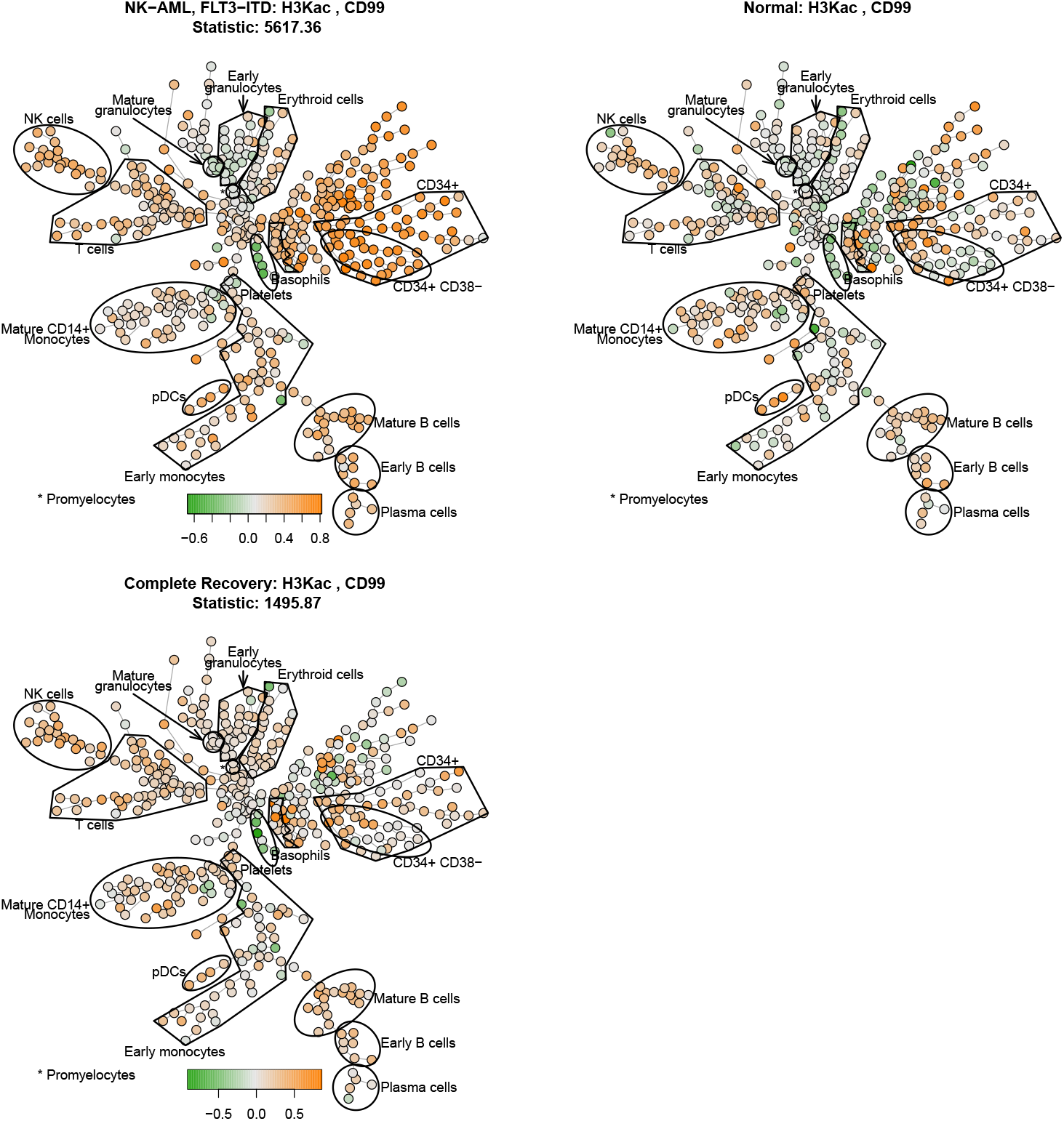
Minimum spanning trees colored by correlation coefficients between histone 3 lysine acetylation (H3Kac) and the CD99 antigen on a relative scale. Top: Correlations computed from pooled data from normal karyotype with a FLT3-ITD mutation (left) compared to pooled data from normal controls (right). Key visual differences can be seen in CD34+ immunophenotype and CD34+ CD38-immunophenotype and in many other nodes. Bottom: Correlations computed from pooled data from patients in the patients without AML after chemotherapy. Key visual differences can be seen in many of the unnannotated other nodes compared to the normals above, but less visual difference between the CD34+ CD38-immunophenotype. Overall there appears to be a decrease in correlation with the patients without AML after chemotherapy compared to the samples from healthy donors.

### Differential correlation between CD99 and H3Kac in CD34+ CD38-cells

We illustrate our approach with the biomarker pair H3Kac and CD99. This pair was found to be differentially correlated for every group of AML samples when compared to normal control samples, but the immunophenotype nodules most contributing to the global *χ*^*δ*^ statistic value varied drastically for each of the groups (Table 5). For this pair each group’s global statistic value was primarily driven by correlation at nodes from the other cells, but the variable contributions of other nodules highlights the advantage and necessity of considering local *χ*^*δ*^ statistics.

**Table 5.**
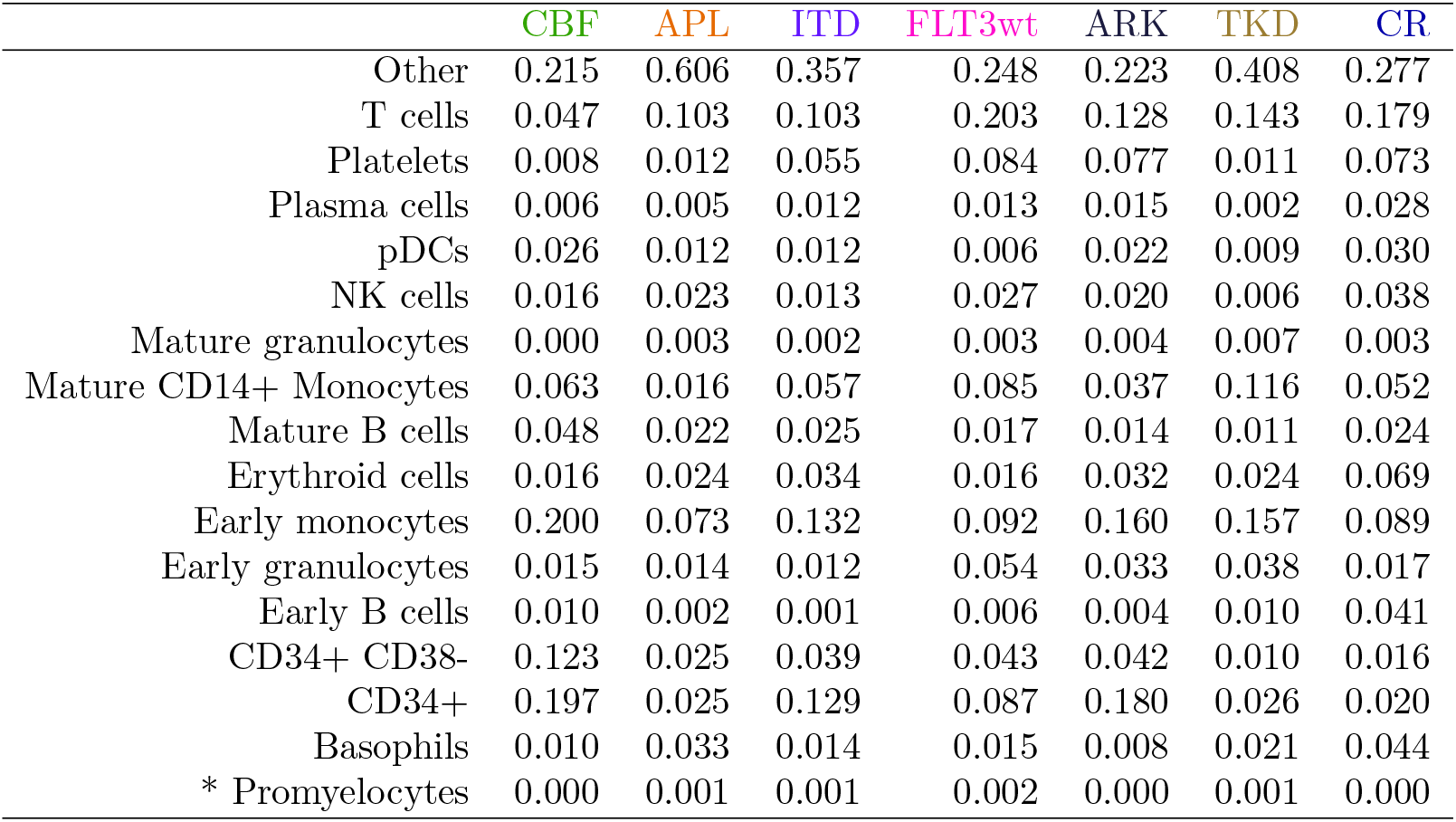
Local *χ*^*δ*^statistic contributions from each immunophenotype nodule when comparing each subtype to the samples from healthy donors. The local *χ*^*δ*^ statistic is the proportion of the global *χ*^*δ*^ statistic that arises from any specified subset of nodes. represent the restriction of the global *χ*^*δ*^statistic to nodes for the CD34+ CD38-nodule. this marker pair was found to be globally differentially correlated for each subtype, but the local contributions from the immunophenotype nodulesdiffers for each comparison.

|  | CBF | APL | ITD | FLT3wt | ARK | TKD | CR |
| --- | --- | --- | --- | --- | --- | --- | --- |
| Other | 0.215 | 0.606 | 0.357 | 0.248 | 0.223 | 0.408 | 0.277 |
| T cells | 0.047 | 0.103 | 0.103 | 0.203 | 0.128 | 0.143 | 0.179 |
| Platelets | 0.008 | 0.012 | 0.055 | 0.084 | 0.077 | 0.011 | 0.073 |
| Plasma cells | 0.006 | 0.005 | 0.012 | 0.013 | 0.015 | 0.002 | 0.028 |
| pDCs | 0.026 | 0.012 | 0.012 | 0.006 | 0.022 | 0.009 | 0.030 |
| NK cells | 0.016 | 0.023 | 0.013 | 0.027 | 0.020 | 0.006 | 0.038 |
| Mature granulocytes | 0.000 | 0.003 | 0.002 | 0.003 | 0.004 | 0.007 | 0.003 |
| Mature CD14+ Monocytes | 0.063 | 0.016 | 0.057 | 0.085 | 0.037 | 0.116 | 0.052 |
| Mature B cells | 0.048 | 0.022 | 0.025 | 0.017 | 0.014 | 0.011 | 0.024 |
| Erythroid cells | 0.016 | 0.024 | 0.034 | 0.016 | 0.032 | 0.024 | 0.069 |
| Early monocytes | 0.200 | 0.073 | 0.132 | 0.092 | 0.160 | 0.157 | 0.089 |
| Early granulocytes | 0.015 | 0.014 | 0.012 | 0.054 | 0.033 | 0.038 | 0.017 |
| Early B cells | 0.010 | 0.002 | 0.001 | 0.006 | 0.004 | 0.010 | 0.041 |
| CD34+ CD38- | 0.123 | 0.025 | 0.039 | 0.043 | 0.042 | 0.010 | 0.016 |
| CD34+ | 0.197 | 0.025 | 0.129 | 0.087 | 0.180 | 0.026 | 0.020 |
| Basophils | 0.010 | 0.033 | 0.014 | 0.015 | 0.008 | 0.021 | 0.044 |
| * Promyelocytes | 0.000 | 0.001 | 0.001 | 0.002 | 0.000 | 0.001 | 0.000 |

To visually depict the variability in nodule contributions, we considered minimum spanning trees for the patients without AML after chemotherapy group and the normal karyotype with a FLT3-ITD mutation group versus the group of normal samples in Fig 5. For the CD34+ CD38-cell populations, we observe a strong visual change in normal karyotype with a FLT3-ITD mutation versus samples from healthy donors and a negligible change when comparing the patients without AML after chemotherapy versus samples from healthy donors. Quantified, the CD34+ CD38-nodule contribution (3.93%) for normal karyotype with a FLT3-ITD mutation is 2.5 times larger than that of the patients without AML after chemotherapy local *χ*^*δ*^ statistic (1.57%).

The correlation between H3Kac and CD99 is present at the level of single cells for several groups when considering biaxial plots in Fig 6. A total regression line, which minimizes both horizontal and vertical displacements between data and the determined line, is added to each subplot in orange to show the overall correlation of the data cloud. The markers are strongly correlated within the CD34+ CD38-particularly in the case of core-binding factor AML; where the global statistic is driven 21.48% by the other nodule and 12.31% by CD34+ CD38-nodule. We note that the correlation is not present for patients without AML after chemotherapy nor samples from healthy donors.

**Fig 6.**
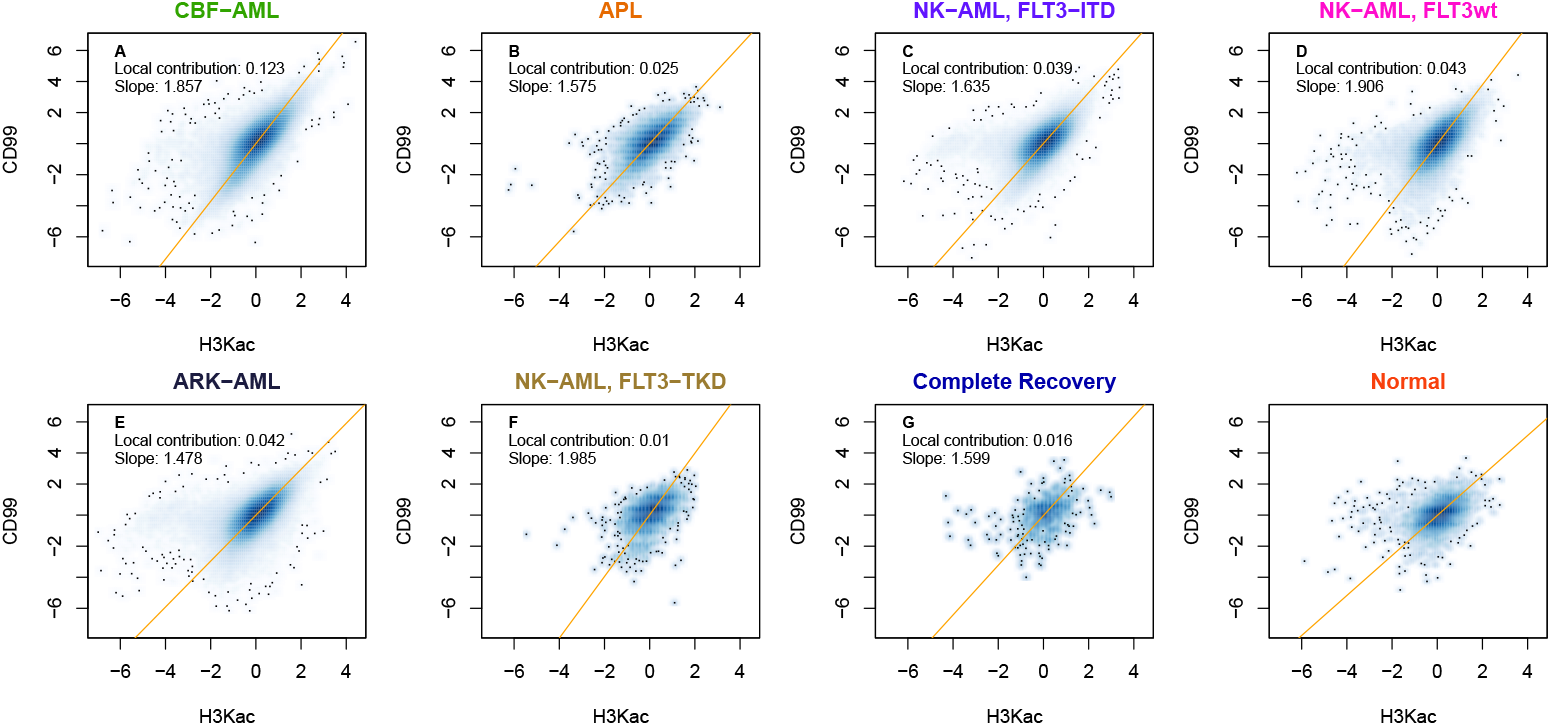
Biaxial plots of histone 3 lysine acetylation (H3Kac) expression versus CD99 antigen expression for individual cells in the CD34+ CD38-immunophenotype nodule. The local statistic contributions from Table 5 and slopes of the orange best fit line of the data cloud have been supplied. The biomarker pair was found to be globally differentially correlated for all sample group comparisons to the normal controls, but for this nodule strong positive correlation is only observed for the samples in Panels A-E. The correlation is not seen in Panels F or G, the sample groups for which the local statistic contribution are smallest, nor in Panel H for the normal controls.

To quantify the visual differences observed in Fig 5 we consider the distribution of correlation coefficients in Fig 7. We observed a marked difference in the shape of the distributions, but also an increase in the average correlation when comparing normal karyotype with a FLT3-ITD mutation (*µ*_*FLT*3−*ITD*_ = 0.6234) versus patients without AML after chemotherapy (*µ*_*Recovery*_ = 0.1976) versus samples from healthy donors(*µ*_*Normal*_ = 0.1978). We also note the similarity in the shape of distributions between patients without AML after chemotherapy and samples from healthy donors.

**Fig 7.**
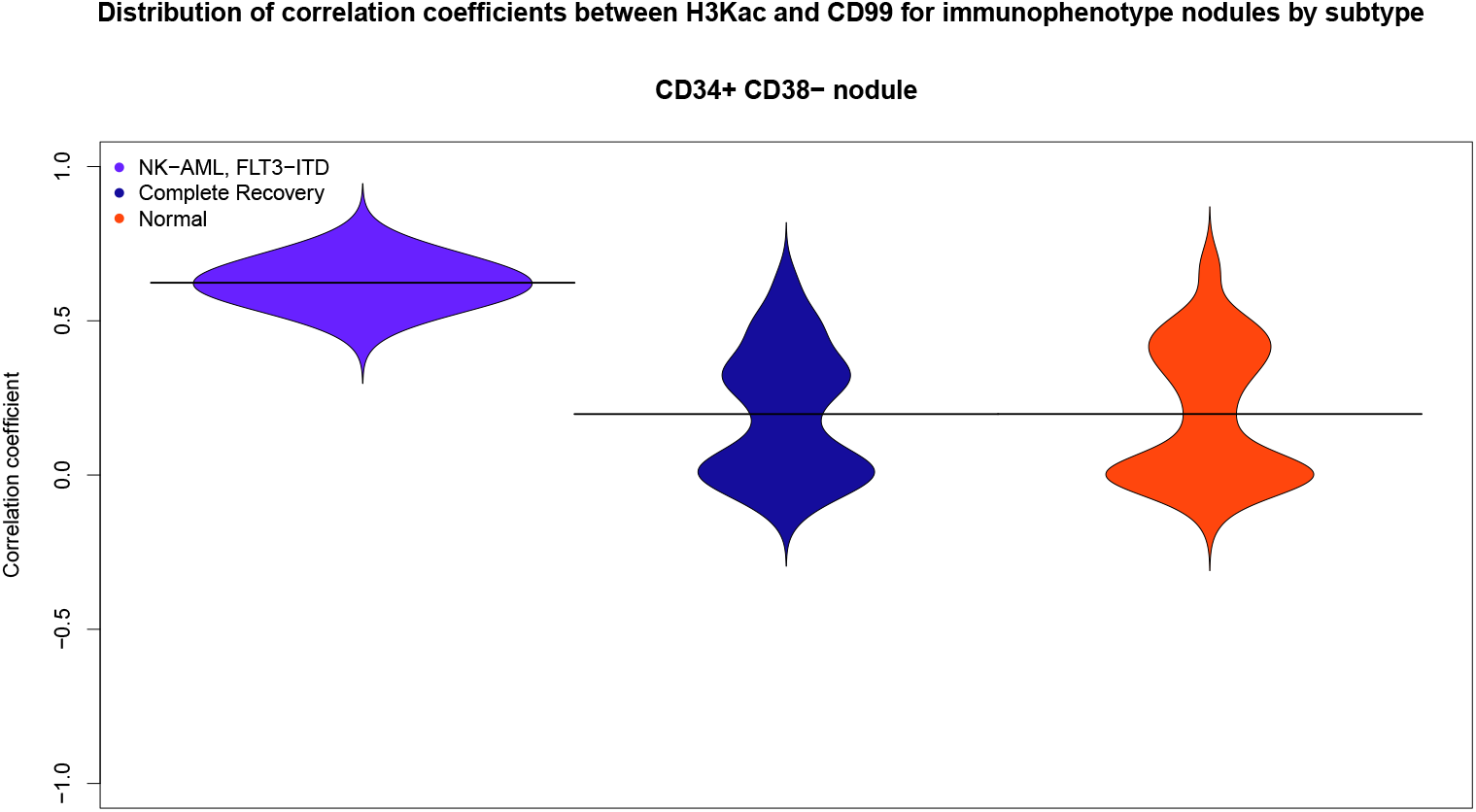
Bean plots depicting the distributions of correlation coefficients between histone 3 lysine acetylation (H3Kac) and the CD99 antigen at each node within the CD34+ CD38-nodule for three sample groups. The average correlation coefficient for each group are shown by the line through its beans. Comparing the distribution for the normal karyotype with a FLT3-ITD mutation samples to the distribution for the samples from healthy donors controls further illustrates the visual difference observed atop the minimum spanning trees in Fig. 5.

## Discussion

Our consideration provides an approach for both testing statistical differences between collections of correlation coefficients and detecting high dimensional events within single cell data. The *χ*^*δ*^ statistic presented is a global measure of differences between collections of correlation coefficients and, when there is additional structure to classify the observations in the collection, a local statistic can be derived by restricting to any subcollection defined by that additional structure. The high-dimensional cytometry data considered here has multiple levels of structure, and restricting the *χ*^*δ*^ statistic to subsets of the data allows for the detection of potentially novel and relevant interactions between quantities measured in the cytometry panel.

More concretely, here the structure provided by surface marker expression leads to a phenotypic classification of the individual cells, and so the *χ*^*δ*^ statistic can be localized to nodules corresponding to particular cell phenotypes. Restricting the statistic to different nodules has proven to be a powerful tool for discerning whether the global measure of differentially correlated proteins are driven by changes in a specific phenotype or driven by many phenotypes.

Localizing the statistic affords an opportunity for correlations, and changes in correlation, to be studied within more nuanced contexts that would not be possible when only computing an aggregate measure. We began with visualizing correlations atop the minimum spanning tree, a step towards visualizing multiple protein interactions as the tree nodes are most often colored proportional by the expression of one protein of interest at a time. For the pair of interest, the visual phenomena atop the trees was explored locally within each nodule. This exploration was accomplished by considering both biaxial plots of the biomarker expressions, allowing us to see whether correlations are present at the single-cell level, and the distributions of their correlation coefficients, allowing quantification of differences in nodule correlation.

The distributional comparisons shown in Fig 7 are similar to methods from Behbehani *et al*. [3] that characterized AML subtypes via marker aberrance, and the average correlations for these distributions help to quantify the visual differences observed in Fig 5. The bimodal characteristic of the distributions for the samples from healthy donors and samples from patients without AML after chemotherapyare interesting to note. While the distribution for the normal karyotype with a FLT3-ITD mutation samples is tightly centered (*σ*_*FLT*3−*ITD*_ = 0.06287872) about the average correlation. The standard deviations for the other distributions are each more than 3 times *σ*_*FLT*3−*ITD*_. Marker pairs that are differentially correlated for AML samples, but not in healthy nor recovered samples can be important for gaining insight into drivers of the disease. Localizations can thus yield testable hypotheses on significant protein interactions.

Localization is also useful before attempting to draw insight from differential correlation. The marker pair H3Kac and CD99 was found to be differentially correlated for all sample groups globally, but when restricting to the CD34+ CD38-nodule it was not the case for two groups: normal karyotype with FLT3-TKD mutation and patients without AML after chemotherapy. These groups are shown in Panels F or G of Fig 6. The marker expressions there, along with the the normal controls in Panel H, do not show strongly correlation as those in Panels A-E. We note also that these groups had the smallest local *χ*^*δ*^ contribution from the CD34+ CD38- (Table 5)..

The presence of the correlations at single cell level for groups like normal karyotype with a FLT3-ITD mutationand the absence for patients without AML after chemotherapyand samples from healthy donorsaligns with the averages of the correlation coefficient distributions seen in Fig 7. This leads us to conclude the statistic is detecting strong correlation and points to the necessity of considering context before reporting differential correlation results. Reporting context is important as differential correlation for this nodule points to an increase of correlation, but when looking at the basophils nodule contributions are different and there is a visual increase in negative correlation when comparing normal karyotype with a FLT3-ITD mutation versus samples from healthy donors.

Another strength of our approach is the statistical method presented is agnostic to the clustering method used to define the subcollections of cells. Thus, we are not restricted to using SPADE before computing correlation coefficients and applying our method, but could also use VISNE [9] with binning or any other contemporary clustering approach for cytometry data.

In many of the sample groups we found that there was a strongly positive correlation in the case of H3Kac and CD99 and the CD34+ CD38-cells had a large contribution to the global statistic value here. When localizing to the CD34+ CD38-nodule, the biaxial plots (Fig 6) confirmed the strong positive correlation at the single cell level between H3Kac and CD99 that was observed visually in the minimum spanning trees for many of the AML subtypes, as well the lack of correlation in the samples from healthy donors and patients without AML after chemotherapy groups.

While correlation does not imply causation, localization can help to explore contexts where it might make sense to begin to consider casual relationships. The strength of the correlation between H3Kac and CD99 is interesting due to CD99’s role in diagnosis and histone 3 being disregulated in many cancers [22]. Determining correlations in context of specific nodules could help in better understanding histone 3 regulation in the context of leukemia, where modifications to histone acetylation are not yet well understood [20]. Moreover, the role of CD34+ CD38-in leukemia diagnosis and outcome [23, 24] makes the presence of this relationship within that nodule interesting to consider.

Correlation is the most common measure of similarity in high-throughput gene expression and large scale omics studies [5, 6], so it is natural to seek a notion of testing correlation when considering entire high dimensional cytometry datasets. Understanding differential correlations between pairs of proteins, like H3Kac and CD99, is a step towards improving the understanding of networks around biomarkers of interest. The localization feature helps discern exactly how correlation relationship present in different nodules.

## Supplement

### Processing clustered data

The acute myeloid leukemia (AML) data was processed in R by four scripts. The first script set up file paths; gathered and stored information about the markers studied (Fig 1) and AML subtypes (Table 1); and created keys on how to match subtypes with samples. This information was then stored them in a R data (RDA) file that is called in all the subsequent scripts. The second script clustered the data in Flow Cytometry Standard (FCS) format with SPADE. SPADE outputs FCS files appended with node and density information as well as layout and Geographical Markup Language (GML) files to recreate the minimum spanning tree with the library iGraph.

The third script separates each appended FCS file into RDA files with cell information for the 483 nodes. For both the cell signaling and the cell cycle panel, the cell per node information is stored for each sample and with all samples pooled by node and samples in each subtype pooled by nodes. To account for differences in cell size within and between samples, as measured by the expression level of an irridium based cell marker (IBC), we removed the effect of cell size to each of the features through a linear regression before computing Pearson correlations.

The fourth script estimated correlation matrices for the 18 markers to be analyzed at each of the nodes through a Bayesian approach [16]. The estimation circumvents issues that arise when a node is empty or contains only a few cells and attempts to correlate data with built-in R functions. When there are no cells (*N* = 0) in a given node, the method estimates the 18×18 correlation matrix *C*_*N*_ for that node to be the identity matrix *I*_18_. Assuming a Wishart prior distribution, the correlation matrix *C*_*N*_ were estimated as follows when there are *N* cells in a node:

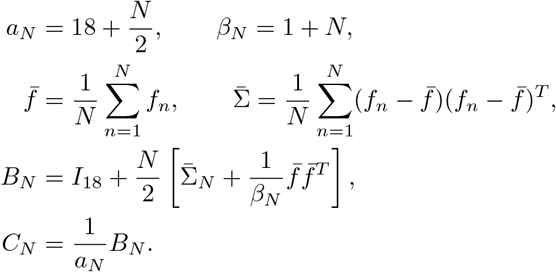

Thus, when there are many cells in a node, then the data-derived estimate overwhelms the prior *I*_18_. Correlation matrices are estimated at every node, and stored in the folders named according to that node, for individual samples, the all-sample pool, and each subtype pool.

### Derivation of the *χ*^*δ*^ statistic

We considered how Pearson correlation coefficients between pairs (153 total) of the 18 markers in the panel reserved for analysis. Given a marker pair *x* and *y*, there is a correlation coefficient at each of the 483 nodes in each group of samples. This resulted in 153 vectors 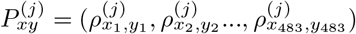, where *j* is a group pooled by sample subtype and each component is a correlation coefficient between markers *x* and *y* in the *i*^*th*^ node determined by SPADE. In order to find statistically significant changes between the group of normal control samples and groups of samples with different subtypes of AML, we sought a distribution on the correlation coefficients and created a test statistic *χ*^*δ*^ (Eq 2) related to a Chi Square, *χ*^2^, statistic.

Our null hypothesis was that all of the correlations arise from the same distribution and thus their distribution parameters were equal. A Fisher z-transform 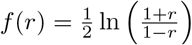 was applied componentwise to 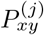 for each *j*. This transform maps the components from a probability distributiJon on [−1, 1] to a normal distribution with 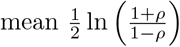 and standard deviation 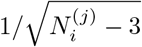, where *ρ* is the true correlation coefficient and 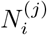 is the number of cells used to estimate 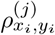. When considering the sum of the 483 squared standardized variables

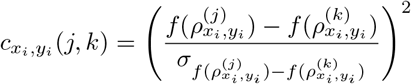

we determined that our statistic for significant aggregate differences in correlation should, in theory, arise from a Chi Square distribution with 483 degrees of freedom, denoted 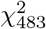.

Due to the number of cells in certain nodes being in the thousands, the effect size at those nodes needed to be taken into consideration in order to accurately determine statistical significance with a 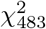 distribution. To account for the effect sizes we regularized our statistic by adding an exchangeability factor *s*_0_ to the standard deviations at each node; an idea borrowed from Tusher *et al*. [17]. We sought a value *s*_0_ that was uniform for all sample comparisons for either cell signaling or cell cycle data. To achieve this we generated values 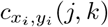 from samples in all groups *j* and *k* at each node *i* for all biomarker pairs *x* and *y*. For all of these choices of *i, j, k, x*, and *y* we sought to find *s*_0_ so that

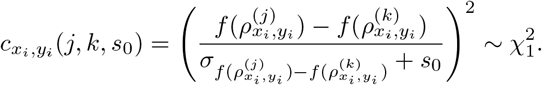

Since a Chi square distribution is uniquely determined by its mean, we arrived at our choice of *s*_0_ by minimizing, over the interval [0,2], the absolute deviation between 1 and the mean of 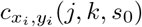 for all *i, j, k, x*, and *y*. The upper bound on the interval [0,2] arose from the case when nodes are empty.

The values of the determined exchangeability factors is found in Table 6. Intuitively, the addition of the *s*_0_ term serves to prevent oversensitivity at nodes with a large number of cells and makes the statistic more conservative overall. Thus, with a uniform value of *s*_0_ in place for each dataset in consideration we define our test statistic to be

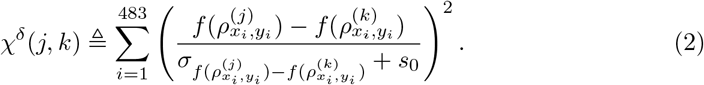

**Table 6.** Exchangeability factors to minimize the difference between the means of the summands 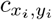 and 1, the mean of a 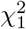. The minimization was performed with the one dimensional optimization function `optimize` in R.

| Cell cycle $s_0$ | Signaling $s_0$ |
| --- | --- |
| 0.02188310 | 0.02092582 |

### Determining the null distribution empirically

Despite including an exchangeability factor *s*_0_, the variability in the number of cells per node within the mass cytometry data led to a discrepancy between the expected and the observed values of the *χ*^*δ*^ statistic. To proceed with hypothesis testing, we chose to define an empirical distribution using the *χ*^*δ*^ statistic to compare the fourteen normal samples. The motivating idea was that an empirical distribution defined from normal comparisons would provide a baseline for significant differences one could expect when comparing vectors of cytometric correlations and anything beyond this baseline difference could confidently be considered significant.

The 44 samples from leukemia samples were non-uniformly distributed to groups corresponding to the seven AML subtypes and we sought to compare these to the group of fourteen normal controls. We investigated the independence to grouping size to avoid introducing false positives due to differences in sizes of groups being compared to normals. Showing independence to group size would have required that the empirical distribution not change when comparing different subgroupings of normal samples. We found that the distribution of *χ*^*δ*^ values produced from the 91 one vs. one comparisons of normal controls (Fig 10) bounded the values produced from the comparisons of *m* vs. *n* subgroupings (Fig 11), where *m* + *n* = 14; we will refer to the latter as complementary subgroupings. The comparisons of the subgroupings were each rerun 30 times. Despite the lack of independence, using the distribution from one vs. one comparisons was a conservative and reasonable choice of distribution since significant findings under it would be significant in all of the distributions from the complementary subgroupings (Fig 11).

**Fig 8.**
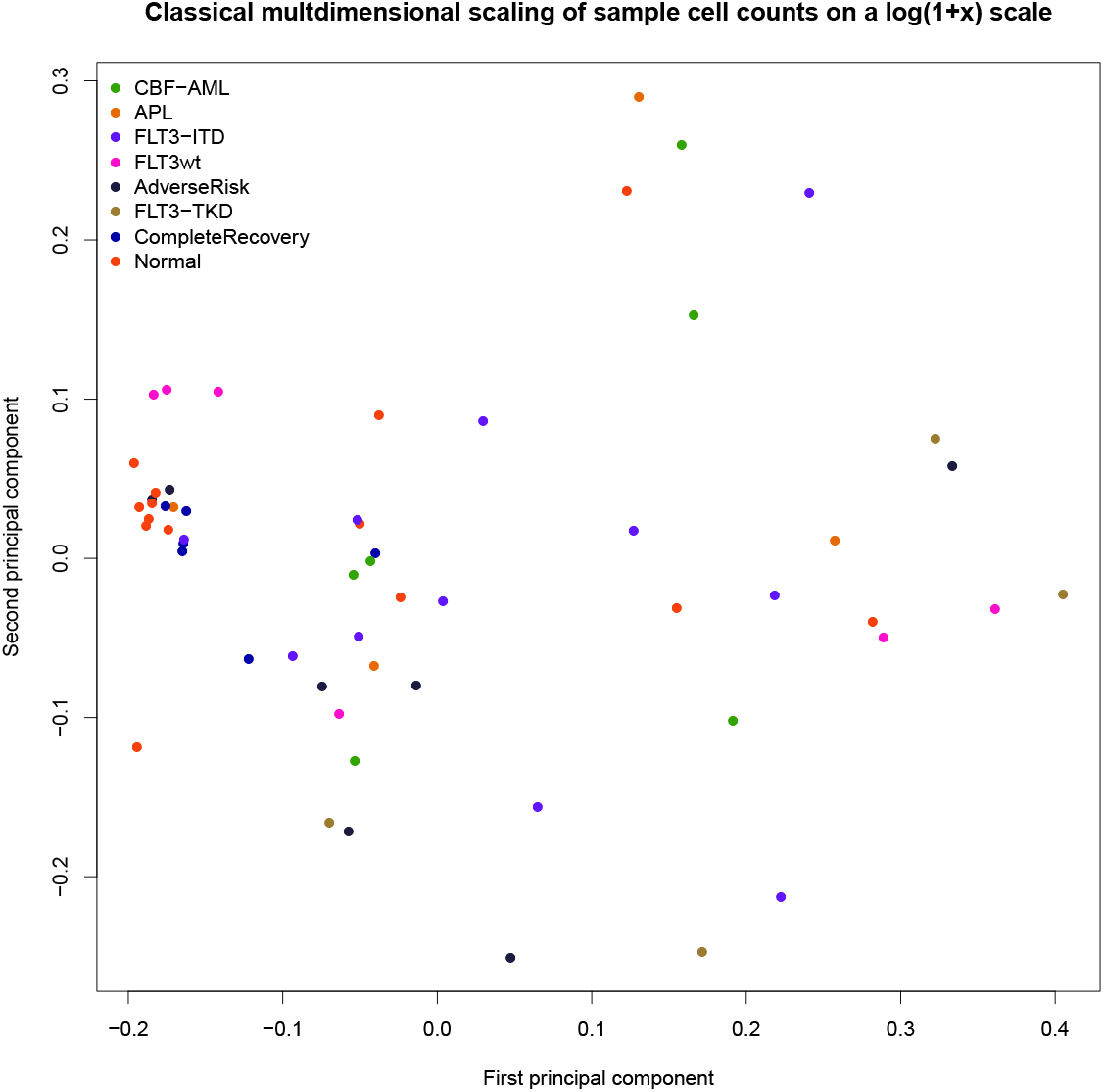
Plot of the cell counts per node for the 58 samples projected onto the first two dimensions determined by classical multidimensional scaling using the correlation distance metric. Cells in each sample were assigned to one of 483 nodes by SPADE [11] and the projection gives a two dimensional summary of the variability of how cells were allocated to the nodes. The cell counts per node are presented on the 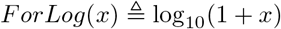 scale. Samples are colored by the disease subtype. The correlation distance was chosen as the metric in the classical multidimensional scaling to avoid the first principal component being driven by cell count.

**Fig 9.**
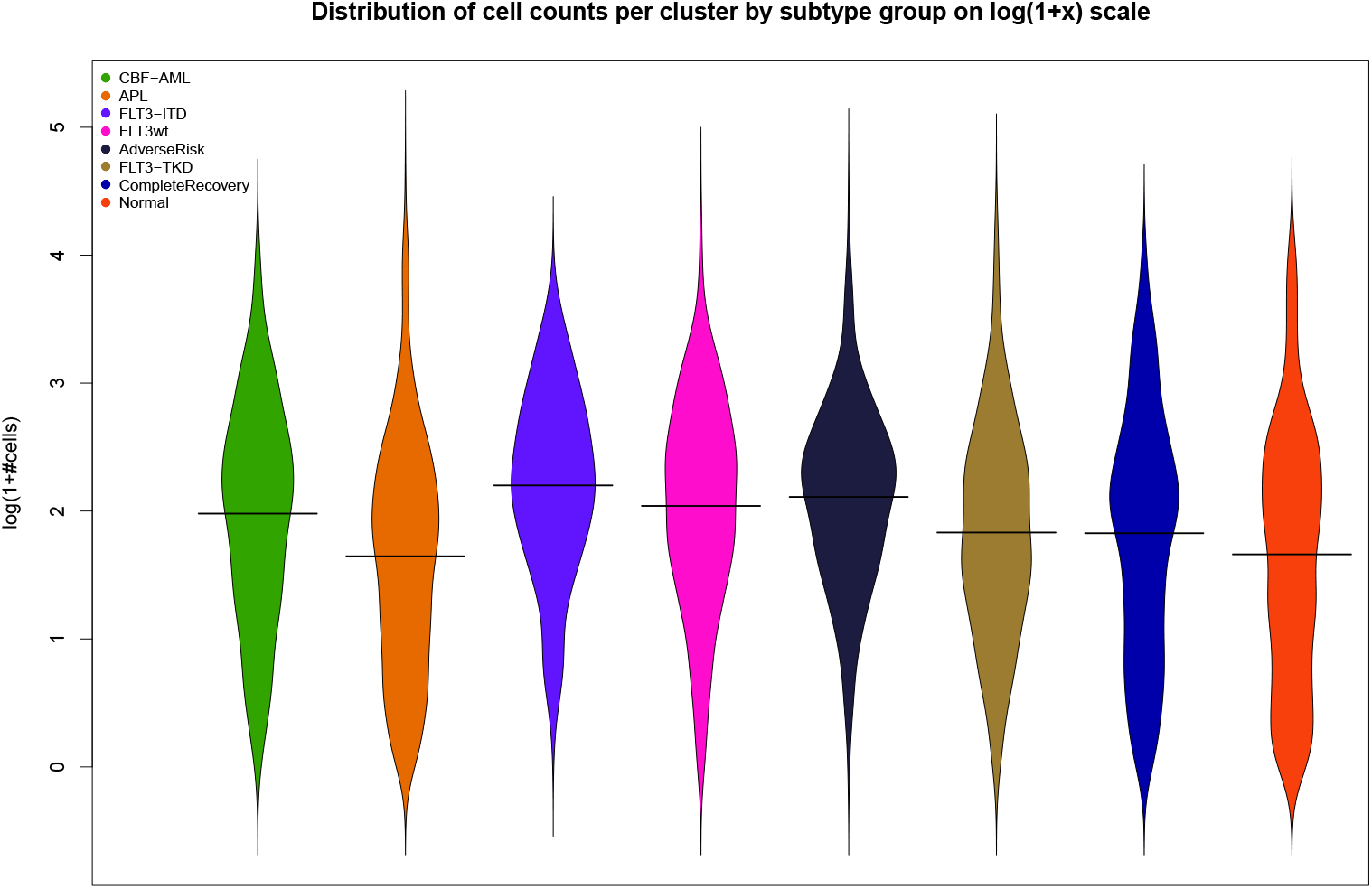
Bean plots for the distribution of cell counts per node separated and colored by subtype. The cell counts per node are presented on the 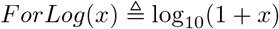 scale. Visually there is a clear pairwise difference of mean and variance for the subtypes. The number cells per node was normalized by the number of cells in the group and adjusted by the average number of cells per sample, The normalization and adjustment was to remove the effect of having a nonuniform number of samples per group on the means.

**Fig 10.**
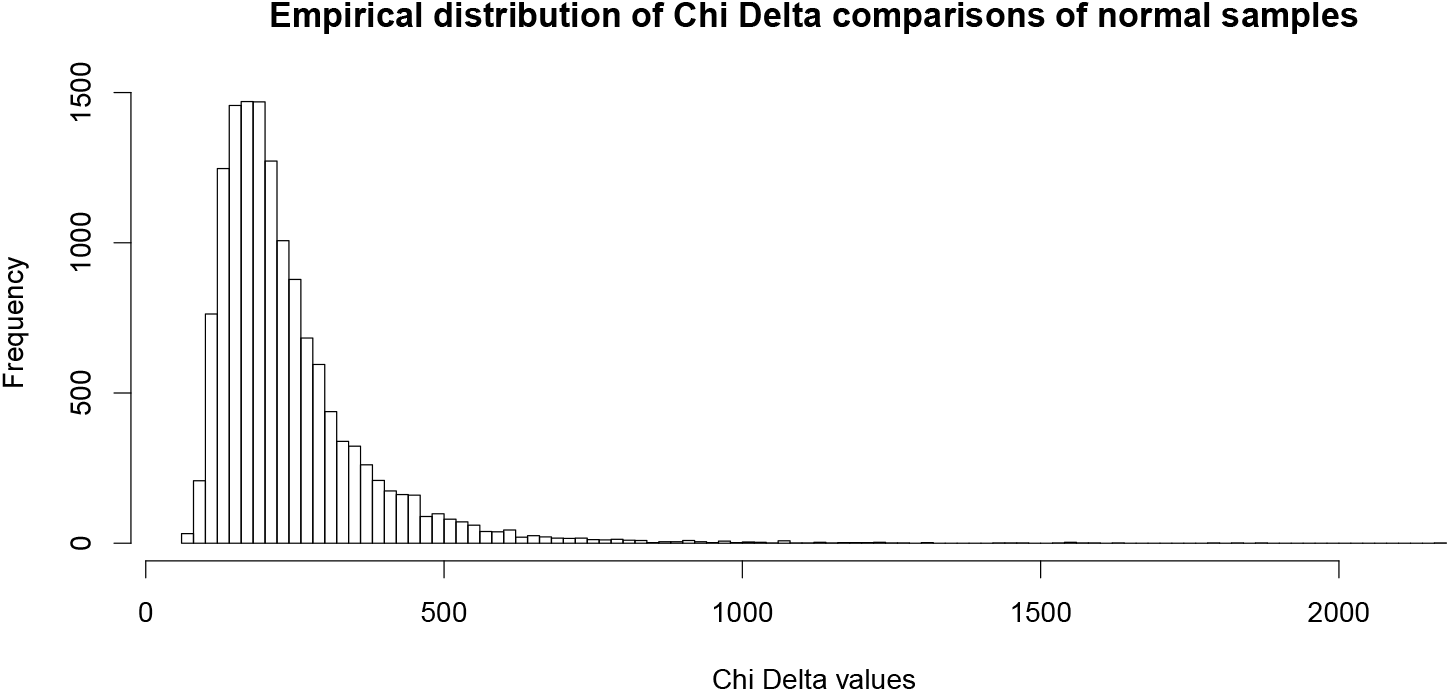
An empirically derived distribution for *χ*^*δ*^ comparisons for the 91 possible pairs normal samples in the case of the cell signaling panel. We show in Fig 11 that this empirical distribution of individual *χ*^*δ*^ comparisons of samples will bound the *χ*^*δ*^ distribution of all complementary groupings. Moreover, the empirical distribution will serve as the null distribution in subsequent statistical comparisons instead of the theoretical distribution. An empirical distribution for the cell cycle data was also derived, but is not shown since analysis for that data is not presented.

**Fig 11.**
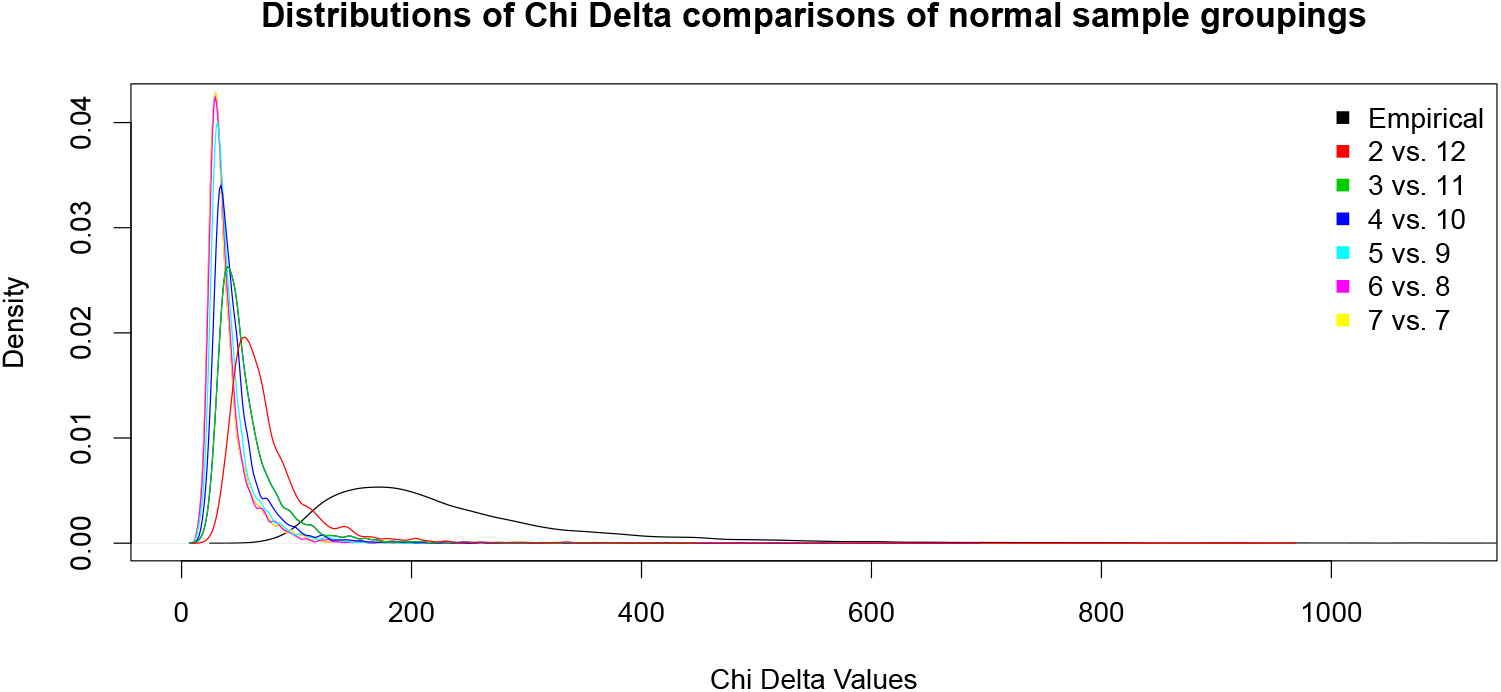
Densities of *χ*^*δ*^ comparisons resulting from complementary groupings of normal samples. Each division of the normal samples into groups of size *m* and *n*, where *m* + *n* = 14 and *m, n ≥* 2, were repeated thirty times. The discrepancy between the subgrouping distributions meant that there the *χ*^*δ*^ statistic was sensitive to the size of groups being compared, but the empirical distribution described in Fig 10 bounded each of the subgrouping distributions. This bound will allow us to circumvent a lack of independence to grouping size. Finally, the bound further ensures the empirical distribution gives a conservative measure of statistical significance.

Using the empirical distribution, we considered nominal *p* values of 0.1, 0.05, 0.01, and 0.001. When considering the nominal *p* value of 0.001 we found the false discovery rate (FDR) value of 0.117 and 0.030 to be satisfactory for the cell cycle and cell signaling datasets, respectively. A summary of the number of significant findings and FDR values at each nominal p value can be seen in Table 3. In the analysis, we choose the level *p* = 0.001 for significance. All marker comparisons found to be significant are listed in Table 7

**Table 7.** Summary of all differentially correlated biomarker pairs and the sample groups for which the pairs were found to be globally differentially correlated compared to normal control samples.

|  | CBF | APL | ITD | FLT3wt | ARK | TKD | CR |
| --- | --- | --- | --- | --- | --- | --- | --- |
| pS6;pATM | FALSE | FALSE | FALSE | TRUE | FALSE | FALSE | FALSE |
| pS6;H3Kac | FALSE | FALSE | TRUE | TRUE | FALSE | FALSE | FALSE |
| pS6;pMAPK | FALSE | FALSE | TRUE | TRUE | TRUE | FALSE | FALSE |
| pS6;CD99 | FALSE | FALSE | FALSE | FALSE | FALSE | TRUE | FALSE |
| pS6;BCat | FALSE | FALSE | FALSE | TRUE | FALSE | FALSE | FALSE |
| pATM;H3Kac | TRUE | FALSE | TRUE | TRUE | TRUE | TRUE | TRUE |
| pATM;pMAPK | FALSE | FALSE | FALSE | TRUE | TRUE | FALSE | FALSE |
| pATM;STAT5 | FALSE | FALSE | TRUE | TRUE | FALSE | FALSE | FALSE |
| pATM;Ki-67 | FALSE | TRUE | TRUE | TRUE | TRUE | TRUE | FALSE |
| pATM;CD99 | TRUE | FALSE | TRUE | FALSE | TRUE | FALSE | FALSE |
| pATM;pErk | FALSE | FALSE | FALSE | TRUE | FALSE | FALSE | FALSE |
| pATM;BCat | FALSE | FALSE | FALSE | TRUE | TRUE | FALSE | FALSE |
| pATM;4EBP1 | FALSE | FALSE | TRUE | TRUE | FALSE | FALSE | FALSE |
| pATM;cPARP | TRUE | FALSE | TRUE | TRUE | TRUE | TRUE | TRUE |
| pATM;pCreb | FALSE | FALSE | TRUE | TRUE | TRUE | FALSE | FALSE |
| H3Kac;pMAPK | TRUE | FALSE | TRUE | TRUE | TRUE | FALSE | FALSE |
| H3Kac;STAT5 | FALSE | FALSE | TRUE | FALSE | FALSE | FALSE | FALSE |
| H3Kac;Ki-67 | TRUE | TRUE | TRUE | TRUE | TRUE | TRUE | TRUE |
| H3Kac;pNFkB | FALSE | FALSE | TRUE | FALSE | FALSE | FALSE | FALSE |
| H3Kac;CD99 | TRUE | TRUE | TRUE | TRUE | TRUE | TRUE | TRUE |
| H3Kac;BCat | TRUE | FALSE | TRUE | FALSE | TRUE | FALSE | FALSE |
| H3Kac;4EBP1 | TRUE | FALSE | TRUE | TRUE | TRUE | TRUE | FALSE |
| H3Kac;pCreb | TRUE | FALSE | TRUE | TRUE | TRUE | TRUE | FALSE |
| H3Kac;DNA-2 | TRUE | TRUE | TRUE | TRUE | TRUE | TRUE | TRUE |
| pMAPK;STAT5 | FALSE | FALSE | FALSE | TRUE | FALSE | FALSE | FALSE |
| pMAPK;Ki-67 | TRUE | FALSE | FALSE | TRUE | TRUE | FALSE | FALSE |
| pMAPK;CD99 | TRUE | FALSE | TRUE | TRUE | TRUE | FALSE | FALSE |
| pMAPK;pErk | TRUE | FALSE | FALSE | FALSE | FALSE | FALSE | FALSE |
| pMAPK;BCat | TRUE | FALSE | FALSE | TRUE | TRUE | FALSE | FALSE |
| pMAPK;pCreb | FALSE | TRUE | TRUE | FALSE | TRUE | FALSE | FALSE |
| STAT3;STAT5 | FALSE | FALSE | FALSE | TRUE | FALSE | FALSE | FALSE |
| STAT3;CD99 | TRUE | FALSE | TRUE | FALSE | FALSE | FALSE | FALSE |
| STAT5;CD99 | FALSE | FALSE | TRUE | FALSE | FALSE | FALSE | FALSE |
| Ki-67;CD99 | TRUE | TRUE | TRUE | TRUE | TRUE | TRUE | FALSE |
| Ki-67;BCat | TRUE | FALSE | FALSE | FALSE | FALSE | FALSE | FALSE |
| Ki-67;4EBP1 | TRUE | FALSE | TRUE | FALSE | TRUE | FALSE | FALSE |
| Ki-67;pCreb | TRUE | FALSE | TRUE | TRUE | FALSE | FALSE | TRUE |
| pNFkB;CD99 | FALSE | FALSE | TRUE | FALSE | FALSE | FALSE | FALSE |
| pNFkB;DNA-2 | FALSE | FALSE | TRUE | FALSE | FALSE | FALSE | FALSE |
| CD99;pErk | TRUE | FALSE | TRUE | FALSE | FALSE | TRUE | FALSE |
| CD99;BCat | FALSE | FALSE | TRUE | TRUE | FALSE | FALSE | FALSE |
| CD99;4EBP1 | TRUE | TRUE | TRUE | TRUE | TRUE | TRUE | FALSE |
| CD99;pCreb | TRUE | TRUE | TRUE | TRUE | TRUE | TRUE | FALSE |
| CD99;DNA-2 | TRUE | TRUE | TRUE | TRUE | TRUE | TRUE | FALSE |
| pErk;4EBP1 | TRUE | FALSE | FALSE | FALSE | FALSE | FALSE | FALSE |
| pErk;pCreb | TRUE | FALSE | FALSE | FALSE | FALSE | FALSE | FALSE |
| pErk;DNA-2 | FALSE | FALSE | TRUE | FALSE | FALSE | FALSE | FALSE |
| cPARP;pCreb | FALSE | FALSE | TRUE | FALSE | FALSE | FALSE | FALSE |
| pCreb;DNA-2 | FALSE | FALSE | TRUE | FALSE | FALSE | FALSE | FALSE |

## References

1. Bendall SC, Simonds EF, Qiu P, El-ad DA, Krutzik PO, Finck R, et al. Single-cell mass cytometry of differential immune and drug responses across a human hematopoietic continuum. Science. 2011;332(6030):687–696.

2. Howlander N. SEER Cancer Statistics Review 1975-2014. http://seercancergov/csr/19752014/. 2017;.

3. Behbehani GK, Samusik N, Bjornson ZB, Fantl WJ, Medeiros BC, Nolan GP. Mass cytometric functional profiling of acute myeloid leukemia defines cell cycle and immunophenotypic properties that correlate with known responses to therapy. Cancer discovery. 2015; p. CD–15.

4. Boyd AL, Aslostovar L, Reid J, Ye W, Tanasijevic B, Porras DP, et al. identification of chemotherapy-induced leukemic-regenerating cells reveals a transient vulnerability of human AML recurrence. Cancer cell. 2018;34(3):483–498.

5. Weinstein JN, Myers TG, O’connor PM, Friend SH, Fornace AJ, Kohn KW, et al. An information-intensive approach to the molecular pharmacology of cancer. Science. 1997;275(5298):343–349.

6. Eisen MB, Spellman PT, Brown PO, Botstein D. Cluster analysis and display of genome-wide expression patterns. Proceedings of the National Academy of Sciences. 1998;95(25):14863–14868.

7. Tenenbaum JB, De Silva V, Langford JC. A global geometric framework for nonlinear dimensionality reduction. science. 2000;290(5500):2319–2323.

8. Roweis ST, Saul LK. Nonlinear dimensionality reduction by locally linear embedding. science. 2000;290(5500):2323–2326.

9. Amir EaD, Davis KL, Tadmor MD, Simonds EF, Levine JH, Bendall SC, et al. viSNE enables visualization of high dimensional single-cell data and reveals phenotypic heterogeneity of leukemia. Nature biotechnology. 2013;31(6):545–552.

10. Sugar IP, Sealfon SC. Misty Mountain clustering: application to fast unsupervised flow cytometry gating. BMC bioinformatics. 2010;11(1):502.

11. Qiu P, Simonds EF, Bendall SC, Gibbs Jr KD, Bruggner RV, Linderman MD, et al. Extracting a cellular hierarchy from high-dimensional cytometry data with SPADE. Nature biotechnology. 2011;29(10):886–891.

12. Shekhar K, Brodin P, Davis MM, Chakraborty AK. Automatic classification of cellular expression by nonlinear stochastic embedding (ACCENSE). Proceedings of the National Academy of Sciences. 2014;111(1):202–207.

13. Platon L, Pejoski D, Gautreau G, Targat B, Le Grand R, Beignon AS, et al. A computational approach for phenotypic comparisons of cell populations in high-dimensional cytometry data. Methods. 2018;132:66–75.

14. Lun AT, Richard AC, Marioni JC. Testing for differential abundance in mass cytometry data. nature methods. 2017;14(7):707.

15. Coombes K, Brock G, Abrams ZB, Abruzzo LV. Polychrome: Creating and Assessing Qualitative Palettes With Many Colors. bioRxiv. 2018; p. 303883.

16. Penny W. Bayesian Inference for the Multivariate Normal. 2014;.

17. Tusher VG, Tibshirani R, Chu G. Significance analysis of microarrays applied to the ionizing radiation response. Proceedings of the National Academy of Sciences. 2001;98(9):5116–5121.

18. Dworzak M, Fröschl G, Printz D, De Zen L, Gaipa G, Ratei R, et al. CD99 expression in T-lineage ALL: implications for flow cytometric detection of minimal residual disease. Leukemia. 2004;18(4):703.

19. Chung SS, Eng WS, Hu W, Khalaj M, Garrett-Bakelman FE, Tavakkoli M, et al. CD99 is a therapeutic target on disease stem cells in myeloid malignancies. Science translational medicine. 2017;9(374):eaaj2025.

20. Agrawal-Singh S, Isken F, Agelopoulos K, Klein HU, Thoennissen NH, Koehler G, et al. Genome wide analysis of histone H3 acetylation patterns in AML identifies PRDX2 as an epigenetically silenced tumor suppressor gene. Blood. 2011; p. blood–2011.

21. Gumy-Pause F, Wacker P, Sappino A. ATM gene and lymphoid malignancies. Leukemia. 2004;18(2):238.

22. Audia JE, Campbell RM. Histone modifications and cancer. Cold Spring Harbor perspectives in biology. 2016;8(4):a019521.

23. Hanekamp D, Cloos J, Schuurhuis GJ. Leukemic stem cells: identification and clinical application. International journal of hematology. 2017;105(5):549–557.

24. Zeijlemaker W, Grob T, Meijer R, Hanekamp D, Kelder A, Carbaat-Ham JC, et al. CD34+ CD38-leukemic stem cell frequency to predict outcome in acute myeloid leukemia. Leukemia. 2019;33(5):1102–1112.

25. Kauffman SA, et al. The origins of order: Self-organization and selection in evolution. Oxford University Press, USA;1993.

26. Levy DE, Darnell J. Stats: transcriptional control and biological impact. Nature reviews Molecular cell biology. 2002;3(9):651–662.

27. Orlova A, Wagner C, de Araujo ED, Bajusz D, Neubauer HA, Herling M, et al. Direct targeting options for STAT3 and STAT5 in cancer. Cancers. 2019;11(12):1930.

